# Neutrophil derived microvesicles induce endothelial cell dysfunction associated with atherosclerotic plaque erosion

**DOI:** 10.1101/2025.02.19.638859

**Authors:** Reece Dow, Erin Card, Sangam Gurung, Paul C. Evans, Victoria Ridger

## Abstract

Atherosclerosis is a major cause of death globally. It is characterised by the development of fibro-fatty lesions in the artery wall that can impede blood flow and lead to myocardial infarction. Historically, research has focused primarily on plaque rupture. However, the importance of erosion of the endothelium leading to thrombosis in plaques with lower lipid content, fewer inflammatory cells and a thick fibrous cap has emerged. Neutrophils have recently been implicated in plaque erosion through the induction of endothelial cell dysfunction. Neutrophils produce 0.1-1μm extracellular vesicles (microvesicles) from their cell membrane that are linked with atherosclerotic plaque progression. The hypothesis that neutrophil microvesicles affect plaque erosion through driving endothelial cell dysfunction and platelet adhesion was investigated. Peripheral blood neutrophils and platelets were isolated from healthy subjects and neutrophils stimulated with native LDL to induce microvesicle release. These microvesicles were found to contain proteases capable of degrading extracellular matrix proteins. Incubation of human coronary artery endothelial cells with neutrophil microvesicles lead to an increase in apoptosis and detachment, and a decrease in migration and proliferation in the endothelial cells. Moreover, microvesicles induced an increase in both platelet P-selectin expression and platelet-endothelial cell interaction. This study demonstrates the propensity of neutrophil microvesicles to promote functions within human coronary artery endothelial cells that may predispose the endothelium to erosion and thrombosis and identifies a potential link between a known risk factor for atherosclerosis, elevated LDL cholesterol, and the production of neutrophil microvesicles. These findings highlight the need for further research to better understand the effects of neutrophil microvesicles in atherosclerosis.

## Introduction

Atherosclerosis is the underlying cause of heart attack and stroke. Characterised by the formation of plaques in the tunica intima of vessels, atherosclerosis causes the progressive narrowing of the vessel lumen and restriction of blood flow. The most severe symptoms occur due to the formation of thrombi following disruption of atherosclerotic plaques, leading to partial or total blockage of the artery lumen. There are broadly two distinct types of atherosclerotic plaques that result in thrombosis: vulnerable plaques and stable fibrous plaques ^1,2^. Vulnerable plaques are highly inflammatory lesions prone to rupture in a process that exposes highly pro-thrombotic materials to the blood and forms a vessel occluding thrombus. Intact fibrous plaques are less well understood; they are significantly less inflammatory than ruptured plaques and often exhibit a smaller lipid core ^3^. Thrombosis associated with stable plaques is believed to occur through a yet to be fully described mechanism that causes the erosion of the plaque endothelium and the exposure of pro-thrombotic components of the extracellular matrix (ECM). Widespread statin use alongside preventative therapies have improved outcomes for patients with or at high risk of suffering from acute coronary syndrome (ACS) resulting from plaque rupture. However, the risk of cardiovascular events following use of these treatments remains unacceptably high. Erosion of intact fibrous plaques may account for this residual risk. Furthermore, as treatments for plaque rupture continually improve, the proportion of cardiovascular events resulting from erosion is increasing ^4^.

The lack of many strong risk factors for superficial erosion has shifted more importance onto the identification of structural aspects of eroded plaques that may indicate a pathological mechanism. Several key processes have been identified that are thought to be involved in plaque disruption and thrombosis. These include the detachment of endothelial cells (ECs) directly beneath the thrombus, accumulation of specific ECM components, the accumulation of platelets to sites of injury and disturbed blood flow. It is likely that several competing processes determine whether ECs are lost from stable plaques. ECM degradation, EC detachment and apoptosis promote erosion whilst proliferation and migration of ECs compensate and protect the plaque from erosion. These processes are likely to be continual, both in diseased and healthy vessels, and are important in normal haemostasis. However, an imbalance in these processes that favours EC dysfunction will likely shift plaques toward a more erosion-prone phenotype, displaying greater EC loss and fewer protective mechanisms.

Recent research suggests neutrophils contribute to plaque erosion. Quillard *et al.* showed neutrophils and neutrophil extracellular traps (NETs) accumulate at sites of apoptotic ECs present in smooth muscle cell-rich plaques ^5^. Moreover, inhibition of neutrophil recruitment under disturbed flow resulted in decreased endothelial permeability, greater continuity and reduced apoptosis compared with controls ^6^. As well as release of NETs, neutrophils are known to release neutrophil microvesicles (NMVs) and these have recently been shown to play a role in plaque development ^7^. Activated neutrophils generate MVs in response to stimulation by exogenous or host-derived factors and contain active MMP-9 ^8^, a matrix metalloproteinase associated with increased anoikis of ECs. NMVs also express PSGL-1 suggesting the potential for NMVs to bind platelets ^7,9^ and have been reported to express tissue factor ^10^ indicating that NMVs also have the potential to regulate platelet activation.

We investigated whether NMVs degrade the ECM and induce endothelial cell dysfunction, thus contributing to the loss of ECs in plaque erosion. We also determined whether NMVs increase platelet activation and adhesion to ECs, contributing to thrombus formation.

## Methods

### Ethics

For human studies, experiments complied with all ethical regulations and were approved by the University of Sheffield Research Ethics Committee (reference number: 031330). Blood samples from healthy subjects over the age of 18 with no known medical conditions were obtained by venepuncture. Participants provided written informed consent and were informed that they could withdraw their consent at any time.

### NMV isolation

NMV isolation was based on the method of Gomez *et al.* ^7^. In brief, human neutrophils were isolated from peripheral venous blood by density gradient separation. Isolated neutrophils were stimulated with native LDL (nLDL, 50 µg mL^-1^; ThermoFisher Scientific, MA) or media alone for 1 h (37 °C in 5% CO2). Neutrophils and large cell debris were then removed by centrifugation (500 × g for 5 min; 2000 × g for 5 min) and the supernatant collected. To pellet NMVs, the suspension was centrifuged at 20,000 × g for 30 min at 4°C. Control samples were spiked with 20 μL of the supernatant from this step. NMV quantification was performed by flow cytometry (LSRII, Becton Dickinson, UK) using Megamix fluorescent calibration beads (BioCytex, France) according to the manufacturer’s instructions. NMVs were measured by LSRII and data analysed using FlowJo software (v.10.6.2, Becton Dickinson, UK.)

### Nanoparticle Tracking Analysis (NTA)

NMV size distribution was assessed by NTA using a ZetaView (Particle Metrix, Germany) with 110 nm calibration beads (Thermofisher, UK), a frame rate of 3.75 frames s−1 and shutter speed of 70. For post-acquisition analysis, parameters were set to a minimum brightness of 25 and a minimum and maximum area of 5 and 999 pixels, respectively. Measurements were taken at 11 positions in the cell, with two cycles of each position. Data was then analysed using Particle Metrix software (ZetaView v8.03.08.03). For experiments where fluorescently labelled NMVs were required, PKH26 or PKH67 fluorescent cell linker kits for general cell membrane labelling were used to label the NMVs directly (Sigma, UK) according to the manufacturer’s instructions.

### Transmission Electron Microscopy (TEM)

Thawed NMV pellets were resuspended in 90 µL of sterile filtered PBS and vortexed for 10 s before use. For negative staining, 5 µL of NMV suspension was absorbed onto a glow-discharged thin-film carbon-coated copper grid for 1 min. The grid was blotted, washed with 50 µL of distilled water for 5 s, blotted and washed again. After blotting for a third time, the grid was incubated with 50 µL of 0.75% uranyl formate for 5 s, blotted, then placed in the stain for a further 20 s. The grid was then blotted thoroughly to remove residual moisture. Grids were immediately visualised (magnification ×68,000) using a Tecnai Spirit G2 Transmission Electron Microscope (ThermoFisher Scientific, MA) at an accelerating voltage of 80 kV and micrographs were taken using a Gatan digital camera (Gatan, CA).

### Western blotting

Isolated neutrophils and NMVs were washed, lysed and centrifuged to remove cellular debris. The protein content of the lysates was determined using microBCA assay according to the manufacturer’s instructions (Thermo Fisher Scientific, MA). Neutrophil and NMV lysates containing equal amounts of protein were combined with LDS sample buffer (Thermo Fisher Scientific, MA), boiled for 10 mins, then run on 4–12% Bis-Tris gel (Thermo Fisher Scientific, MA). Separated proteins were transferred onto 0.45 µm PVDF membranes (GE Healthcare, IL). Membranes were blocked and exposed to rabbit anti-CD63 and anti-CD9 antibodies (1:2000 dilution, Abcam, UK) overnight at 4°C. Membranes were washed and goat anti-rabbit Horseradish Peroxidase conjugated secondary antibody was added (1:3000 dilution, Cell Signalling Technology, MA, catalogue number: 7074) for 1 hr at room temperature. Membranes were washed again and incubated for 1 min at room temperature with Amersham ECL Select™ western blotting detection reagent (Cytvia, MA). Membranes were imaged using a ChemiDoc™ MP Imager (Bio-Rad,CA). Images were processed with Image Lab software version 6.1 (Bio-Rad,CA).

### Surface antigen labelling of neutrophil microvesicles and analysis by flow cytometry

Flow cytometry allowed the analysis of NMV surface molecules derived from the parent neutrophil. NMVs produced in response to nLDL were resuspended to 1,000 NMV/μL in 100μL 4°C cell staining buffer (BioLegend, UK) and were incubated with 5μg/mL CD66-FITC antibody on ice for 45 mins (BioLegend, UK). Samples were made up to 200μL with cell staining buffer and centrifuged at 20,000 ×g for 30 minutes at 4°C to pellet the labelled NMVs. The resulting supernatant was removed with care taken to avoid disturbing the pellet. Samples were resuspended in 400μL 4°C cell staining buffer prior to analysis by flow cytometry.

### Quantification of neutrophil microvesicle protease contents using profiler array

Reagents provided with the Proteome profiler human protease array (R&D Systems, USA) were prepared according to the manufacturer’s instructions. PBS and nLDL stimulated NMVs were lysed via freeze/ thaw cycles prior to resuspension in 100 μL PBS and diluted in 1.5 mL 1X array buffer. Nitrocellulose membranes containing 35 separate protease capture antibodies were blocked with 2 mL array buffer in a 4-well multi-dish for 1 h at room temperature on a rocking platform. 15 μL of reconstituted biotinylated protease detection antibody cocktail was added to 1.5 mL NMV samples and incubated at room temperature for 1 h. The blocking array buffer was aspirated from the 4-well multi-dish, 1.5 mL NMV samples added to membranes and incubated with protease detection antibody cocktail overnight at 4 °C on a rocking platform. Membranes were removed and wash buffer added for 10 min on a rocking platform. This process was repeated three times. Membranes were incubated with 2 mL of streptavidin-horse radish peroxidase (HRP) for 30 min at room temperature. Membranes were washed three times with wash buffer to remove unbound streptavidin-HRP, placed within protective plastic and incubated with chemi reagent mix (provided by manufacturer) for 1 min. The fluorescent signal was quantified through multiple exposure times using a BioRad ChemiDoc XP system. The mean pixel intensity for the negative control spots was quantified and subtracted from the mean pixel intensity of the pair of duplicate spots on the membrane corresponding to each protease. This analysis was performed using ImageJ software (version 1.52).

### MMP-9 and Neutrophil Elastase (NE) ELISA

MMP-9 and NE DuoSet ELISA kits (R&D Systems, USA) were used to measure protein concentration in isolated NMVs in accordance with the manufacturer’s instructions. 96-well ELISA plates were coated with MMP-9 or NE capture antibody overnight. Plates were washed and blocked with 300 μL of 1% BSA and incubated for 1 h at room temperature. NMV pellets were resuspended in 100 µl filtered PBS and lysed by multiple freeze-thaw cycles. 100 μL of diluted standards/samples was added to the appropriate wells. Samples were added to wells and the plates were covered and incubated for 2 h at room temperature. 100 μL of 250 ng mL^-1^ detection antibody was added to each well. The plates were covered and incubated for a further 2 h at room temperature. 100 μL of a 40-fold dilution of streptavidin-HRP was added to each well. The plates were incubated away from direct light for 20 min at room temperature. The aspiration wash step was repeated to remove all unbound streptavidin-HRP. A substrate solution was prepared from a 1:1 ratio of colour reagent A (hydrogen peroxide) and colour reagent B (tetramethylbenzidine). 100 μL of this was added to each well. Plates were incubated for 20 min at room temperature away from direct light. After 20 min 50 μL of stop solution was added to each well to inhibit HRP and induce a colour change from blue to yellow. The optical density of each well was quantified using a VarioSkan microplate reader (ThermoFisher, USA). All wash steps were performed using a Geneflow ELx50 plate washer (Biotek Instruments, USA).

### NE activity

A fluorometric NE activity assay kit (Sigma, UK) was used to assess NE activity in NMVs in accordance with the manufacturer’s instructions. A concentration range of NE standards was included to serve as a reference. NMVs were resuspended to 1×10^4^ NMV μL^-1^ in sterile PBS before being lysed via freeze-thaw method. 10 μL NMV supernatant was diluted by an equal amount to serve as control. Wells were made up to 50 μL with additional NE assay buffer. 20 μL of sample was added to wells and made up to 50 μL with additional NE assay buffer. 50 μL NE substrate mix (48 μL NE assay buffer + 2 μL NE substrate) was added to 50 μL of samples per well to make 100 μL per well. Fluorescence was measured over 20 min at 37 °C (ex = 380nm/ em = 500nm). Two time points were chosen that fell within the linear range and the corresponding fluorescence values recorded. The fluorescence of the background standard was subtracted. NE activity was calculated according to the manufacturer’s instructions.

### Gelatin degradation assay

DQ-gelatin was also used in a microplate assay to assess gelatin degradation by NMVs. DQ-gelatin is FITC-labelled and the fluorescent signal is quenched when DQ-gelatin is not degraded. Following degradation by proteases, a fluorescent signal is emitted proportional to the degree of degradation. This fluorescence can be quantified using a plate reader. 80 μL sterile filtered PBS was added per well of a black 96-well microplate. 20 μL of DQ-gelatin (250 μg mL^-1^) was added per well. Freshly isolated NMVs were resuspended in phenol red-free media. Phenol red-free media containing NMV supernatant was used as a negative control. 100 μL of NMV (1×10^4^ µL^-1^) samples and negative control were added to wells and the microplate incubated in the dark at room temperature. Fluorescence intensity was measured using a microplate reader (ex = 485/ em = 530) and measurements recorded every 15 min for 4 h.

### Human coronary artery endothelial cells

Primary human coronary artery endothelial cells (HCAEC) were purchased from Promocell (Germany). Cells were cryopreserved immediately after isolation, shipped and stored in liquid nitrogen until use. Cells were then defrosted and cultured at 37 °C, 5% CO2 in specialised endothelial growth cell medium (MV2, Promocell, Germany) until confluent. Primary cultures were used for experiments at passage 4–6.

### Endothelial cell detachment

HCAEC detachment was assessed using an assay adapted from previous published studies ^5^. The principal behind this method is that the treatment, in this case NMVs, makes ECs more vulnerable to detachment induced by low levels of trypsin. 3×10^4^ HCAEC were cultured in a 6 well plate on 70 μg/mL collagen type III for 24 h prior to treatment with supernatant control or NMVs (ratio 1:2 HCAEC:NMV) ± 400 ng TIMP-1 for 24 h. HCAECs were washed with sterile PBS and incubated with 6.25μg mL^-1^ trypsin-EDTA for 3 min before being washed with fresh cell culture media (MV2, Promocell). The remaining cells that had adhered to the surface were then imaged by widefield microscopy and detaching cells quantified using ImageJ software (version 1.52). Detaching cells were identified as highly refractive rounded cells.

### Flow cytometry analysis of HCAEC apoptosis

3×10^5^ HCAECs were seeded into a 6 well plate coated with 70 μg mL^-1^ collagen type III and incubated with 300 µL supernatant control or 1:2 ratio of NMVs for 24 h. Following treatment, HCAECs were detached with 25 μg mL^-1^ trypsin-EDTA and resuspended at a concentration of 2×10^5^ HCAECs mL^-1^ in MV2 cell culture media. 1μL CellEvent Caspase-3/7 Green Detection Reagent was added to 1 mL of HCAECs and incubated at 37°C for 30 min protected from light. Apoptosis was analysed by flow cytometry. Fluorescence was excited using a 488 nm laser and emission collected at 530/30 nm. Unlabelled HCAECs and HCAECs treated with 1µM staurosporine (positive control) were used to set the gating strategy, and median fluorescence intensity and percentage positivity values assessed by FlowJo software (version 10.7.1).

### HCAEC proliferation assay

96-well plates were coated with 70 μg mL^-1^ collagen type III. HCAECs were seeded at a low concentration of 3.5 x 10^3^ HCAECs per well to ensure confluency was not reached during the experiment. HCAECs were cultured for 24 h to allow for adherence. HCAECs were treated with 300 µL supernatant control or NMVs (ratio 1:2 HCAEC:NMV) for 24 h. HCAECs were then washed with sterile PBS, fixed with 4% paraformaldehyde for 5 mins, washed again and permeabilised with 0.2% triton-×100 for 15 mins. The washing step was repeated and non-specific binding blocked with 5% FBS for 1 h. HCAECs were washed again and incubated with 1μg/mL Alexa 700 conjugated Ki-67 monoclonal antibody (ThermoFisher, USA) for 2 h at room temperature. HCAECs were washed and counterstained with 300 nM DAPI prior to imaging by widefield fluorescence microscopy (Leica, AF6000).

### Endothelial cell migration assay

HCAECs migration was assessed using time-lapse phase contrast microscopy. Ibidi culture inserts were placed in wells of a 6-well cell culture plate. 3×10^4^ HCAECs were seeded within each quadrant of the cell culture inserts and allowed to adhere for 24 h. The culture media in each quadrant was aspirated and replaced with media containing supernatant control or NMVs (ratio 1:2 HCAEC:NMV) and cultured for 2 h. The cell culture inserts were then removed using sterilised tweezers to create a sterile and reproducible cell gap between confluent HCAEC monolayers. HCAECs were washed with sterile PBS to remove non-adherent HCAECs. NMVs or NMV supernatant was added once again to relevant wells and HCAECs were viewed by time-lapse widefield phase contrast microscopy (Leica, AF6000) at 37°C and 5% CO_2_. Images were taken every 30 min for 24 h. ImageJ software was used to measure the gap/wound area at each time point. The data were then expressed as percentage wound closure.

### Platelet isolation from whole human blood

Venous blood was obtained from healthy subjects (as detailed above) and was centrifuged at 260 x g for 20 min to separate platelet rich plasma (PRP), which was collected and diluted 1:1 with HEP buffer (140nM NaCl, 2.7 mM KCl, 3.8 mM HEPES, 5 mM EGTA) containing 2 μM prostaglandin E1 (PGE1). PRP was mixed gently and centrifuged at 100 x g for 20 min to pellet contaminating red blood cells. The resulting supernatant was removed and centrifuged at 800 x g for 20 mins to pellet platelets. The platelet pellet was washed without resuspension in wash buffer (10 mM sodium citrate, 150 mM NaCl, 1mM EDTA, 1% w/v dextrose). The platelets were resuspended in Tyrode’s buffer (134 mM NaCl, 12 mM NaHCO3, 2.9 mM KCl, 0.34 mM Na2HPO4, 1 mM MgCl2, 10 mM HEPES) containing 5 mM glucose, freshly added BSA (3 mg mL^-1^) and 1 μM PGE1.

### Flow cytometry analysis of P-selectin expression on platelets

Flow cytometry was used to quantify the effect of NMVs on platelet P-selectin expression. 5 μL PRP was added to tubes containing supernatant control or 5 µl NMVs (1 x 10^4^ µl^-1^) ± CD62P-PE (BD Biosciences; clone AC1.2) or CD42a-FITC (BD Biosciences, clone Beb1) and incubated for 20 mins at room temperature. Platelets were then fixed in buffer containing 4% formaldehyde for 10 min and analysed by flow cytometry. Data was analysed using FlowJo software (version 10.7.1).

### Platelet adhesion assay

Widefield fluorescence microscopy was used to assess the effect of NMVs on the interaction between platelets and HCAECs. 2 x 10^4^ HCAEC were seeded into wells of a 48-well plate coated with 70 μg mL^-1^ collagen type III and cultured for 24 h. HCAECs were washed with sterile PBS and treated with 1:5 ratio of HCAEC:NMV for a further 24 h. HCAECs were washed again with sterile PBS. Isolated platelets were labelled with the cell membrane dye PKH-26 (ThermoFisher, USA) and resuspended in 2 mL diluent C. A 4 μM dye solution was prepared by diluting PKH-26 in diluent C and immediately added to the platelet solution to create a final dye concentration of 2 μM. Platelets were incubated at room temperature for 3 min with gentle agitation. The staining reaction was stopped by adding 4 mL of 1% BSA solution and incubating at room temperature for 1 min. Platelets were centrifuged at 800 x *g* for 20 min and washed in fresh MV2 cell culture media. This wash step was repeated twice more. HCAECs were incubated with 1 x 10^6^ PKH-26 labelled platelets per well for 30 min, washed gently three times with sterile PBS, and fresh MV2 cell culture media was added to each well. Cells were immediately imaged by widefield fluorescence microscopy at 37°C and 5% CO_2_ to assess platelet adhesion. Image analysis was performed using Image J (version 1.52).

### Statistical analysis

Results are presented as mean ± SEM throughout. Statistical analysis was performed using GraphPad Prism version 10.2.3 (GraphPad Software, CA). Data was analysed assuming Gaussian distribution using one-way ANOVA followed by paired or unpaired t-test, Tukey’s post hoc test for multiple comparisons or Dunnett’s post hoc test to compare to control values. P values of less than 0.05 were considered significant. For all experiments n numbers relate to different donors for both HCAEC and neutrophils/platelet. In experiments where percentages are shown, data are expressed as the percentage of the mean of the control samples.

## Results

### Native LDL induces microvesicle formation by neutrophils

LDL is a known risk factor in atherosclerosis and, whilst LDL is not found in the same quantities in stable plaques as rupture prone plaques, a large-scale study of patients determined that those who suffered an acute coronary event as a result of plaque erosion had mean serum LDL levels above the healthy range ^3^. We have previously described the characteristics of NMVs derived from fMLP stimulated neutrophils in detail ^7^ but NMVs from nLDL stimulated neutrophils have not been investigated. Stimulation of neutrophils with nLDL resulted in the release of significantly higher numbers of NMVs compared to those released by unstimulated cells (Figure 1A). NMVs released by nLDL stimulated neutrophils were characterised in accordance with the latest ISEV guidance ^11^. TEM demonstrated the size and structure of isolated NMVs (Figure 1B). A representative size frequency distribution histogram of NMVs released in response to nLDL stimulation and a representative frame from videos used to track Brownian motion of nLDL particles is shown in Figure 1C. A cluster of peaks was identified between 105-255 nm and the modal peak was 165 nm. This size range is within the size range of large extracellular vesicles/MVs (100-1,000 nm) ^12^. NMVs were also characterised for their expression of the tetraspanins CD9 and CD63 by western blot analysis and were found to express CD9 but undetectable levels of CD63 (Figure 1D). The neutrophil specific marker, CD66b, was also detected on the surface of NMVs by flow cytometry (Figure 1E).

**Figure 1.**
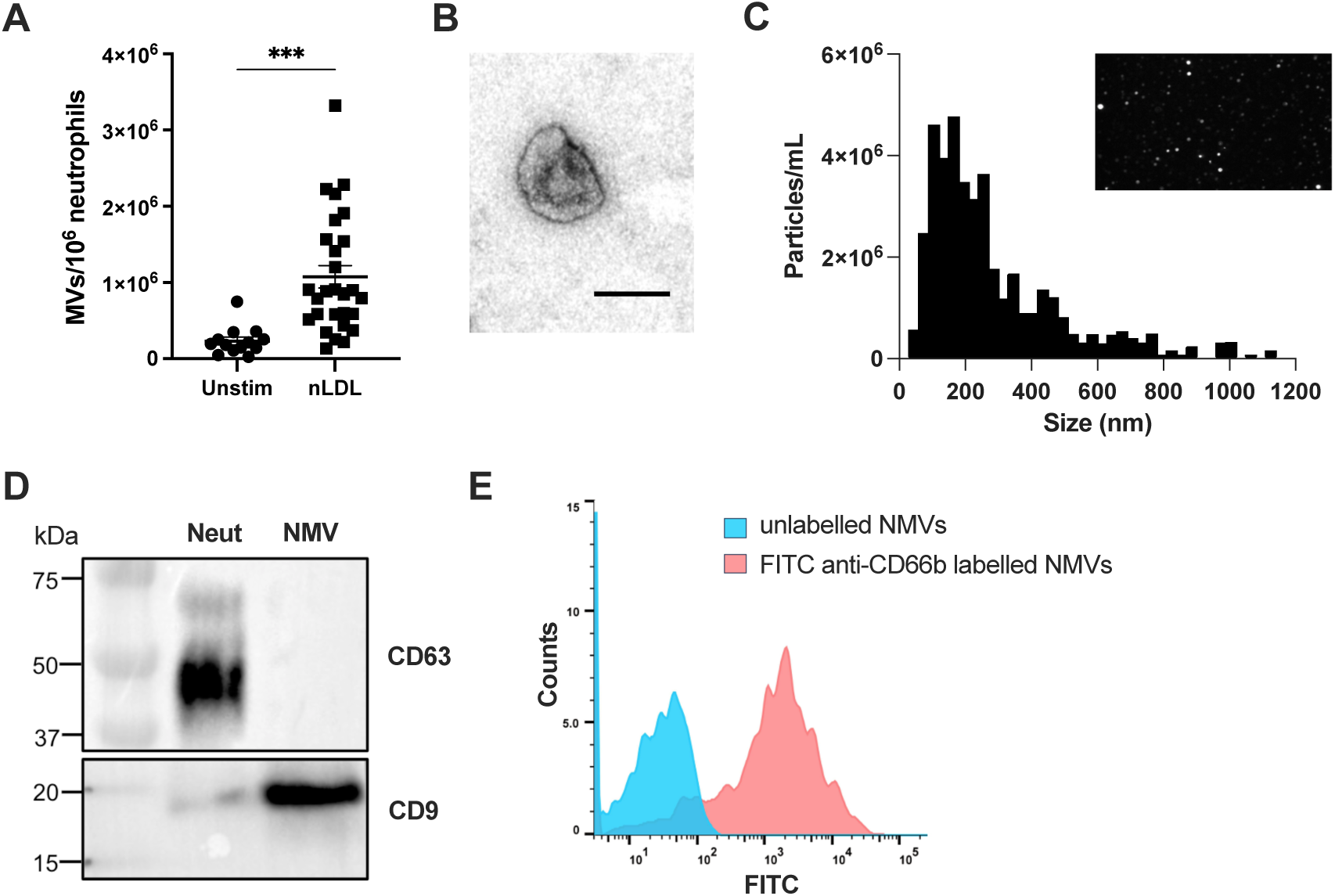
Neutrophil microvesicle biogenesis in response to native low-density lipoprotein stimulation. Human neutrophils were isolated from whole blood of healthy volunteers and resuspended in PBS (unstim) ± native LDL (50 μg/mL; nLDL) for 1 h at 37 °C. NMVs were quantified by flow cytometry **(A).** Data are expressed as mean number of events per 1×10^6^ neutrophils ± SEM, n=13-28. Statistical significance was assessed by one-way ANOVA followed by an unpaired t-test. *** P=<0.001. **(B)** Transmission electron micrograph of a negatively stained NMV sample on a carbon-coated copper grid. Magnification ×68,000, scale bar = 0.2µm. **(C)** ZetaView particle tracking was calibrated with 110nm polystyrene beads prior to size distribution analysis of 200nm, 500nm and 760nm calibration beads. A representative size frequency distribution histogram from 3 separate isolations of NMVs released from nLDL stimulated neutrophil is shown along with a representative frame from videos used to track Brownian motion of nLDL particles. **(D)** Western blot analysis of CD9 and CD63 protein expression in neutrophils (Neut) and NMVs. **(E)** Flow cytometry analysis of CD66b expression on NMVs.

### Neutrophil microvesicles contain active proteases

Having established that nLDL induces NMV formation, we next sought to determine whether these NMVs contained proteases, since neutrophils are known to contain several proteolytic factors that are involved in matrix degradation and inflammation ^13^. To determine which proteases were present in NMVs during neutrophil stimulation, a protease array kit (R&D Systems, USA) was used to screen 35 human proteases for their relative abundance. From 35 screened proteases 9 were detected in NMVs from unstimulated neutrophils and 7 detected in NMVs from nLDL stimulated neutrophils (Figure 2A). Of the different proteases detected, 6 were detected in both NMV populations. A difference was observed in relative quantities of MMPs between the NMV populations (Figure 2A and B). MMP-9 was the most abundant protease detected in NMVs from nLDL stimulated neutrophils followed by MMP-8, whereas this was reversed in NMVs from unstimulated neutrophils with MMP-8 being most abundant, followed by MMP-9. Interestingly, there appeared to be a differential incorporation of 4 proteases in NMVs; Cathepsin V was only detected in NMVs from nLDL stimulated neutrophils and, conversely, Cathepsin X/Z/P, ADAM8 and Urokinase were detected in NMVs from unstimulated neutrophils but were absent in NMVs from nLDL stimulated neutrophils.

**Figure 2.**
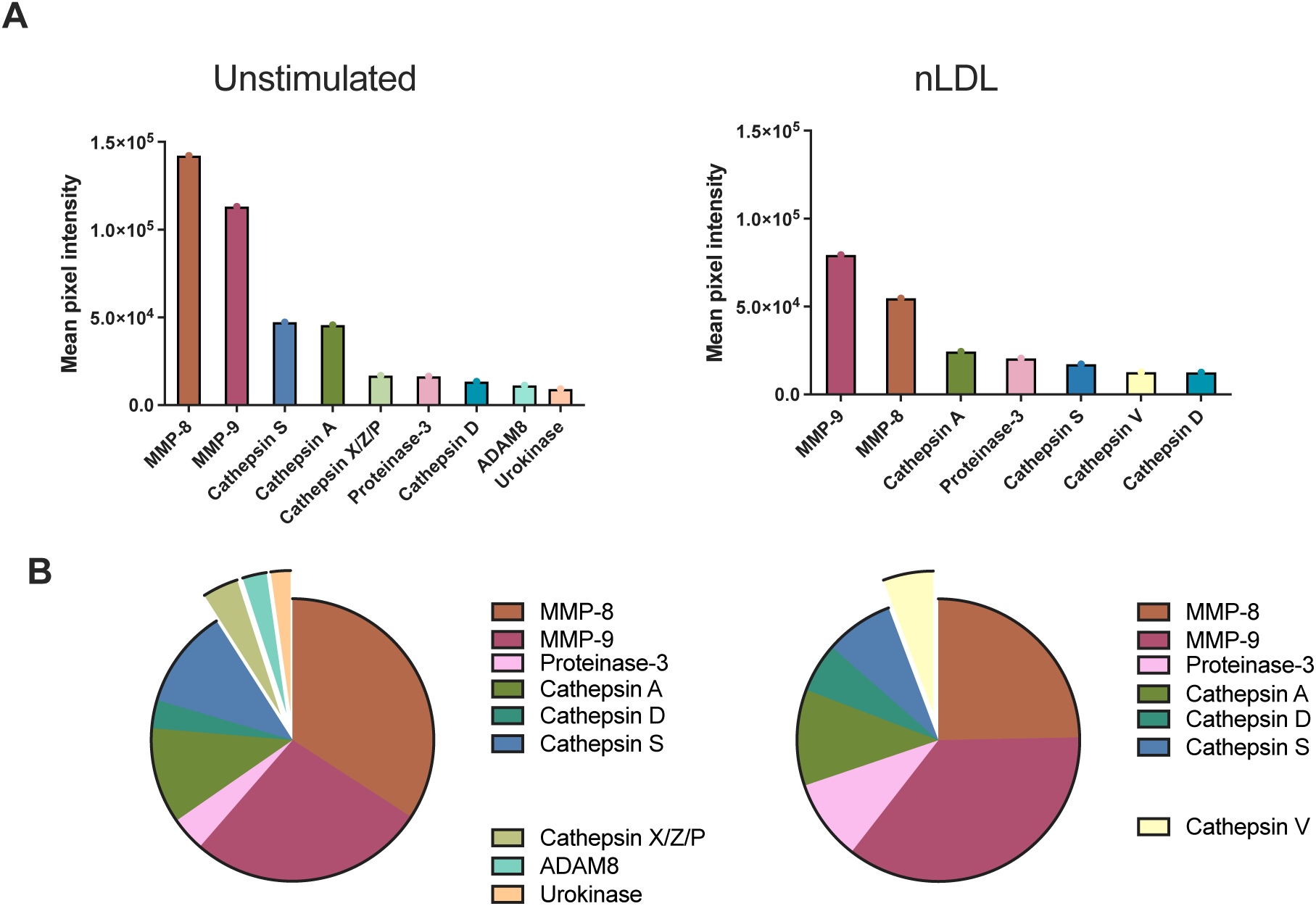
Protease content in neutrophil microvesicles. Neutrophil microvesicle (NMV) protease content was analysed by protease profiler array. NMV lysates were added to nitrocellulose membranes containing 35 separate protease capture antibodies. Following sample incubation and treatment with a detection reagent, protease content was detected by chemiluminescent analysis. The pixel intensity corresponded to the concentration of protease in the sample. **(A)** Relative concentrations of detected proteases in NMVs from unstimulated (unstim) and native LDL stimulated (nLDL) neutrophils. Data are presented as mean pixel intensity from 2 separate arrays. **(B)** Pie charts demonstrating relative levels of protease content in NMVs derived from unstimulated and nLDL stimulated neutrophils. Detached segments represent proteases not conserved between the populations of NMVs.

MMP-9 is a potent protease involved in ECM turnover and has been linked to atherosclerotic plaque rupture ^14^. To further investigate the levels of MMP-9 in NMVs, total MMP-9 concentrations (both active and pro-enzyme forms) in NMVs from unstimulated and nLDL stimulated neutrophils were measured by ELISA. Similar concentrations of MMP-9 were detected per NMV in nLDL stimulated and unstimulated conditions (Figure 3A). However, when MMP-9 concentration was expressed per 1×10^7^ neutrophils to account for the greater number of NMVs released in response to nLDL, a higher level of MMP-9 was detected (32.6 ± 10.5 ng/mL, mean ± SEM, n=4) compared to NMVs from unstimulated neutrophils (4.4 ± 0.2 ng/mL, mean ± SEM, n=3; Figure 3B). No MMP-9 was detected in the supernatant control.

**Figure 3.**
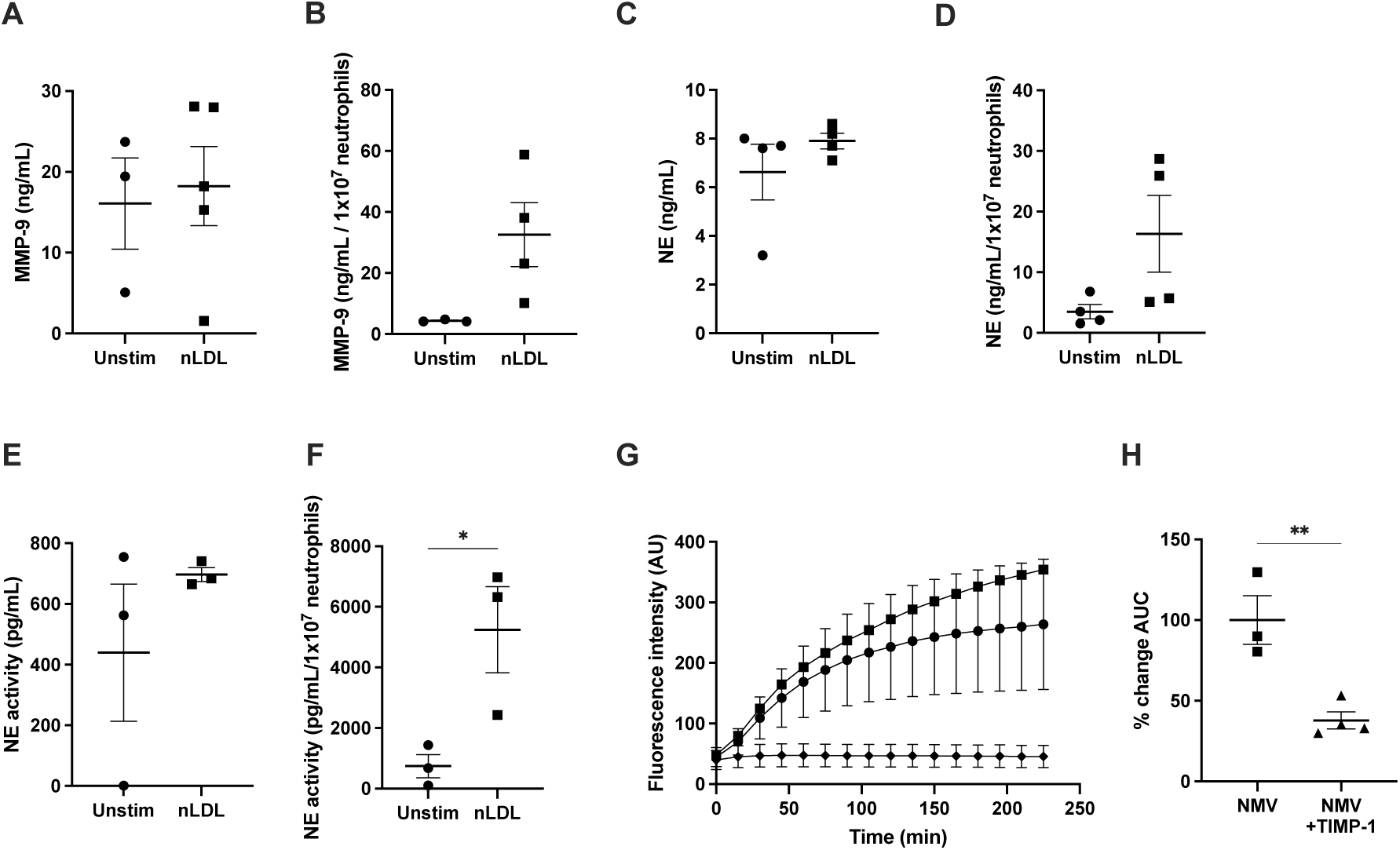
Protease activity in neutrophil microvesicles. MMP-9 and NE concentrations in neutrophil microvesicle (NMV) lysates from unstimulated and nLDL stimulated neutrophils were measured by ELISA **(A and C)** and expressed per 1×10^7^ neutrophils from which NMVs were derived **(B and D)**. NE activity was measured in NMV lysates using an activity assay kit **(E)** and expressed per 1×10^7^ neutrophils from which NMVs were derived **(F)**. To measure protease activity, NMVs from unstimulated (closed circles), native LDL stimulated (nLDL) neutrophils (closed squares), or supernatant control (closed diamonds) were incubated with DQ-gelatin **(G)**. Fluorescence emitted following degradation was quantified using a plate reader and normalised to activity at t=0 for supernatant control. **(H)** The experiment was repeated with NMVs from nLDL stimulated neutrophils ± TIMP-1 (4 µg/mL) and normalised to the level of activity at t=0 for NMVs + TIMP-1. Data expressed as mean ± SEM, n=3-5. Statistical significance was assessed by one way ANOVA followed by an unpaired t-test. * p=<0.05.

NE is another protease released by neutrophils during activation. This protease was not one of those measured in the protease array, so the presence of NE in NMVs was investigated using a standard ELISA. There was no significant difference in NE concentration per NMV between NMVs from unstimulated and nLDL stimulated neutrophils (Figure 3C). When NE concentration was expressed per 1×10^7^ neutrophils, greater NE was detected in the nLDL condition (16.35 ± 6.35 ng/mL, mean ± SEM, n=4) than the unstimulated condition (3.5 ± 1.2 ng/mL, mean ± SEM, n=4; Figure 3D) but this was not statistically significant. NE was not detected in the supernatant control. Having determined the presence of NE in NMVs, we next assessed NE activity in NMVs derived from unstimulated and nLDL stimulated neutrophils. As with total concentrations of NE, there was no significant difference in NE activity per NMV in NMVs from unstimulated compared to nLDL stimulated neutrophils (Figure 3E). However, NE activity was significantly increased in NMVs from nLDL stimulated neutrophils (5243 ±1420 ρg/mL, mean ± SEM, n=3) when compared to NMVs from unstimulated neutrophils (740 ± 385 ρg/mL, mean ± SEM, n=3, p=0.0376) when NE concentrations were expressed per 1×10^7^ neutrophils (Figure 3F). These data suggest that NE is present in NMVs from both unstimulated and nLDL stimulated neutrophils and the increase in the number of NMVs released in response to nLDL results in higher levels of NMV-associated NE activity from these neutrophils.

Healthy endothelial function depends on the close interaction with the ECM, the loss of which can induce a form of apoptosis known as anoikis ^15^. Having shown NMVs contain MMP-9 and neutrophil elastase, which are known to degrade collagens ^16^, we sought to determine whether NMVs had the ability to degrade ECM. NMVs derived from unstimulated and nLDL stimulated neutrophils were incubated with DQ-gelatin (ThermoFisher, USA) which fluoresces following degradation. Fluorescence was measured over time and the area under the curve quantified for statistical analysis (Figure 3G). NMVs from unstimulated and nLDL stimulated neutrophils degraded gelatin whereas supernatant control induced minimal degradation (Figure 3G). Addition of TIMP-1 significantly reduced degradation (62.2% decrease, SEM ± 5.24, n=4, p=0.0068) (Figure 3H). These data indicate that NMVs from both unstimulated and nLDL stimulated neutrophils are capable of degrading analogues of the ECM.

### Neutrophil microvesicles induce detachment and apoptosis of arterial endothelial cell

Following determination that NMVs contain active proteases, we determined whether NMVs induce detachment of HCAECs from the ECM. HCAECs were cultured on collagen III before treatment with NMVs derived from neutrophils treated with nLDL or supernatant control. The percentage of cells detaching was significantly increased in the presence of NMVs (mean ± SEM = 23.8 ± 3% n=7) compared with control (mean ± SEM = 9.1 ± 0.9%, n=7, p=0.0003) (Figure 4A). When TIMP-1 was added, the percentage of detaching HCAECs significantly decreased (mean ± SEM = 10.3 ± 2.4%, n=4, p=0.0052) compared with NMV alone (mean ± SEM = 23.8 ± 3%, n=7) returning the percentage of detaching HCAECs to levels comparable to the supernatant control.

**Figure 4.**
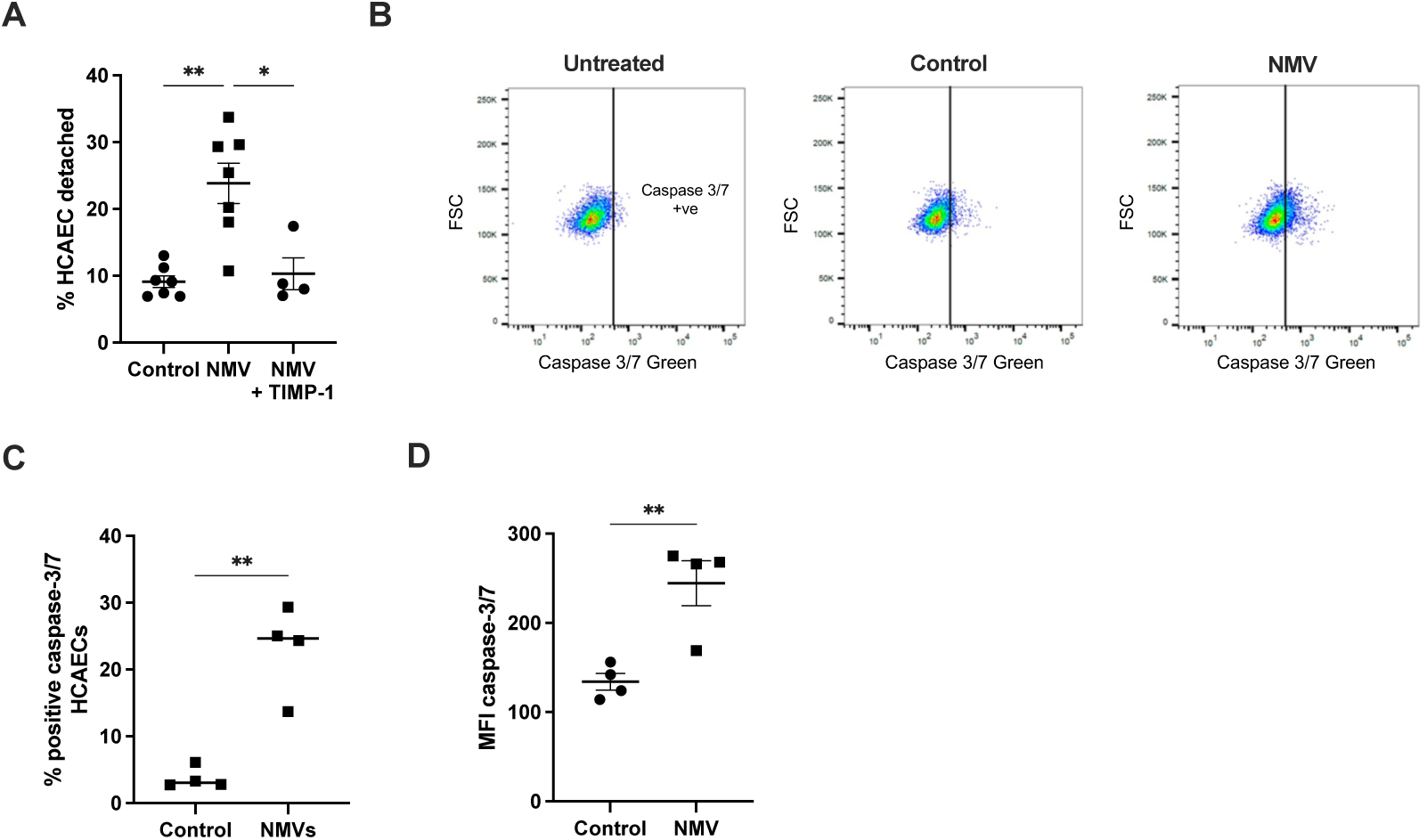
Coronary artery endothelial cell detachment and apoptosis induced by neutrophil microvesicles. **(A)** Human coronary artery endothelial cells (HCAECs) were cultured on 70 μg/mL collagen type III before treatment with supernatant control or NMVs from nLDL stimulated neutrophils (1:2 HCAEC:NMV ratio) ± TIMP-1 (4 µg/mL) for 24 h. HCAECs were then treated with 6.25 μg/mL trypsin-EDTA to induce detachment. Detaching HCAECs were quantified by widefield phase-contrast microscopy and the total number of HCAECs quantified in each condition. To assess apoptosis, HCAECs were cultured for 24 h prior to treatment with supernatant control or NMVs. After 20 h, positive staining for caspase 3/7 was assessed by flow cytometry **(B)** and the percentage **(C)** and mean fluorescence intensity **(D)** of caspase-3/7 positive HCAECs was quantified. Data expressed as mean ±SEM, n= 4-7 (A) n=4 (B). Statistical significance was assessed by one-way ANOVA with Tukey’s multiple comparisons test or paired t-test. * p<0.05, ** p<0.01.

HCAEC viability relies heavily on the interactions with the ECM. As NMVs contain active proteases and can induce HCAEC detachment from a collagen substrate, we investigated whether NMVs increase apoptosis in HCAECs. HCAECs were incubated with NMV supernatant or NMVs from nLDL stimulated neutrophils for 20 h and active caspase-3/7 analysed by flow cytometry. Presence of NMVs significantly increased the percentage of caspase-3/7-positive HCAECs (mean ± SEM = 56.9 ± 2.4, n=4, p=0.0011) compared to supernatant controls (mean ± SEM = 3.75 ± 0.8, n=4; Figure 4B and C) as well as significantly increasing caspase-3/7 mean fluorescence intensity (MFI) (mean ± SEM = 245 ± 25.2, n=4, p=0.0063) compared to supernatant treated controls (mean ± SEM =134 ± 9.3, n=4; Figure 4D).

### Neutrophil microvesicles reduce proliferation and migration in arterial endothelial cells

Whilst processes that contribute to the removal of ECs overlying a fibrous plaque, such as ECM degradation and EC apoptosis, are important in plaque erosion, it is likely that the impairment of processes protective against erosion, such as proliferation and migration, may also contribute. It is likely that ECM degradation, EC detachment and apoptosis promote erosion whilst proliferation and migration of ECs compensate and protect the plaque from erosion. Having determined that NMVs increase HCAEC detachment and apoptosis, the effect of NMVs on HCAEC proliferation was investigated using Ki-67 staining. The number of Ki-67 positive HCAECs significantly decreased in the presence of NMVs (mean decrease = 48.7% ± 4.8, n=4, p=0.0037) compared with supernatant treated control (Figure 5A).

**Figure 5.**
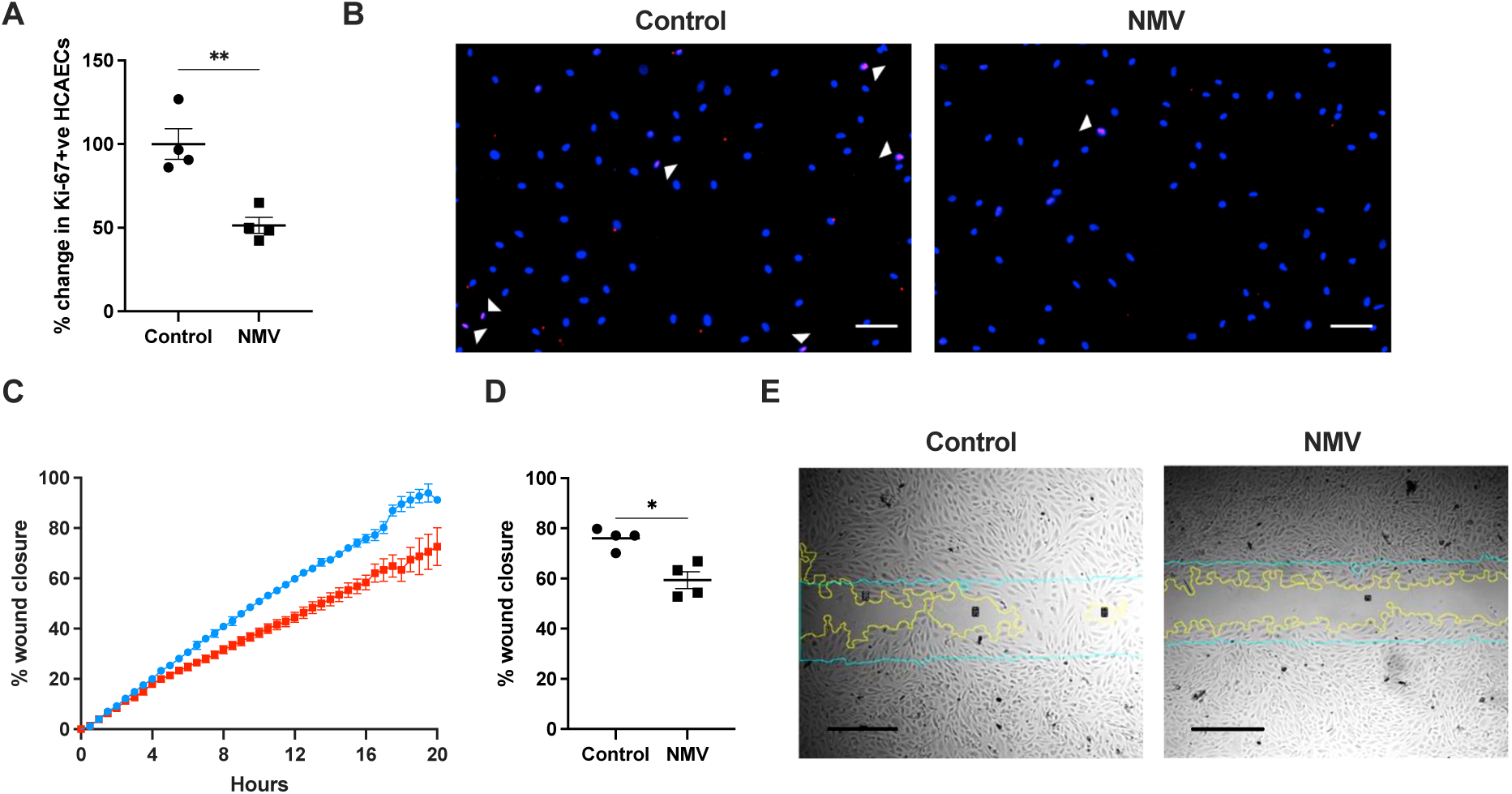
Reduced proliferation and migration in coronary artery endothelial cells in the presence of neutrophil microvesicles. **(A)** Human coronary artery endothelial cells (HCAECs) were cultured on 70 μg/mL collagen III for 24 h prior to treatment with supernatant control or neutrophil microvesicles (NMVs) for a further 24 h. Proliferation was assessed by Ki-67 staining in the nucleus using fluorescence microscopy and presented as the percentage change compared to the mean of the control samples. **(B)** Representative micrographs showing nuclear staining with DAPI (blue), and Ki-67 (red) staining in HCAECs. Ki-67 colocalisation in the nucleus is shown in magenta and highlighted by arrowheads. Scale bar = 100 µm. To assess migration, HCAECs were in cell culture inserts either with supernatant control or neutrophil microvesicles (NMV; 1:2 HCAEC:NMV ratio). Cell inserts were removed to leave a gap in the monolayer and migration of HCAECs was recorded over 20 h by time-lapse phase contrast microscopy. **(C)** Percentage of wound area closed by supernatant treated and NMV-treated HCAECs over 20 h. A time-point consistently within the linear range was chosen (16 h) for direct comparison **(D)**. Representative images of wound healing response by HCAECs 16 h **(E)**. The cyan line indicates the initial wound boundary, and the yellow line represents the edge of the HCAEC monolayer on either side of the wound area. Scale bar = 500 µm. Data expressed as mean ± SEM, n=4. Statistical significance was assessed by paired t-test. * p<0.05, ** p<0.01 Link to video files for wound healing assay: https://drive.google.com/drive/folders/1q4-cQiGQ41rvNDp6MDW32zkviJ0JOSE4?usp=sharing

Following erosion of endothelial cells from the plaque, the underlying pro-thrombotic extracellular matrix is exposed to blood flow, raising the risk of thrombosis. When endothelial cells encounter a ‘wound’ that exposes their underlying matrix, they migrate to the opposing cell front to close the gap. If this ability to recover a wound is impaired, the matrix remains exposed for longer, increasing the risk that a thrombotic event will occur. To assess the effect of NMVs on the ability of HCAECs to migrate, gaps were created in HCAEC monolayers by placing wound healing assay inserts (Ibidi, Germany) into a 6 well plate. HCAECs were then cultured with supernatant control or NMVs and imaged every 30 min by automated time-lapse phase-contrast microscopy over 24 h. Wound recovery was assessed as the percentage of cell gap area closed over time. Initially, the wound recovery response was comparable in both conditions, but from 5 h onwards the rate of wound recovery observed in the two populations of HCAECs began to diverge. Supernatant-treated HCAECs maintained a linear trend whilst the rate of wound recovery in NMV-treated HCAECs began to decrease (Figure 5B). After 16 h NMV treated HCAECs had closed a significantly smaller percentage of the cell gap (mean % EC coverage of gap = 59.34 ± 3.41, n=4, p=0.0058) compared to control (mean % EC coverage of gap = 76.04 ± 2.08, n=4; Figure 5C). 16 h was chosen as the time point for comparison as it consistently fell within the linear phase of wound recovery in supernatant control HCAECs.

### NMVs increase P-selectin expression levels on platelets and platelet adhesion to endothelial cells

P-selectin is stored within α-granules in resting platelets but is rapidly translocated to the surface following platelet activation by stimuli including ADP and thrombin. Membrane expression of P-selectin is a commonly used marker of α-granule exocytosis and platelet activation ^17^. To assess NMV effects on platelet P-selectin expression, ADP was used as a priming stimulus as NMVs alone were insufficient to induce increases in P-selectin expression. Quantification of the median fluorescence intensity (MFI) for P-selectin indicated a modest but significant increase in P-selectin expression following NMV treatment (mean % increase in MFI: 10 ± 1.16, n=3, p=0.025) compared to supernatant control (Figure 6A). Quantification of P-selectin positive platelets indicated no change in the number of platelets that were positive for P-selectin expression following NMV treatment compared with supernatant controls. A high percentage of platelets were P-selectin positive in both NMV treated and supernatant control platelets (Figure 6B). However, this increase in platelet P-selectin expression did not have an effect on platelet aggregation (Figure 6C). The effect of NMVs on platelet adhesion to HCAEC was also investigated (Figure 6D). Analysis of the number of fluorescently-labelled platelets to HCAEC found that increased numbers of platelets were present when HCAEC were pretreated with NMVs (mean % increase = 52.5 ± 15.84, n=6, p=0.015).

**Figure 6.**
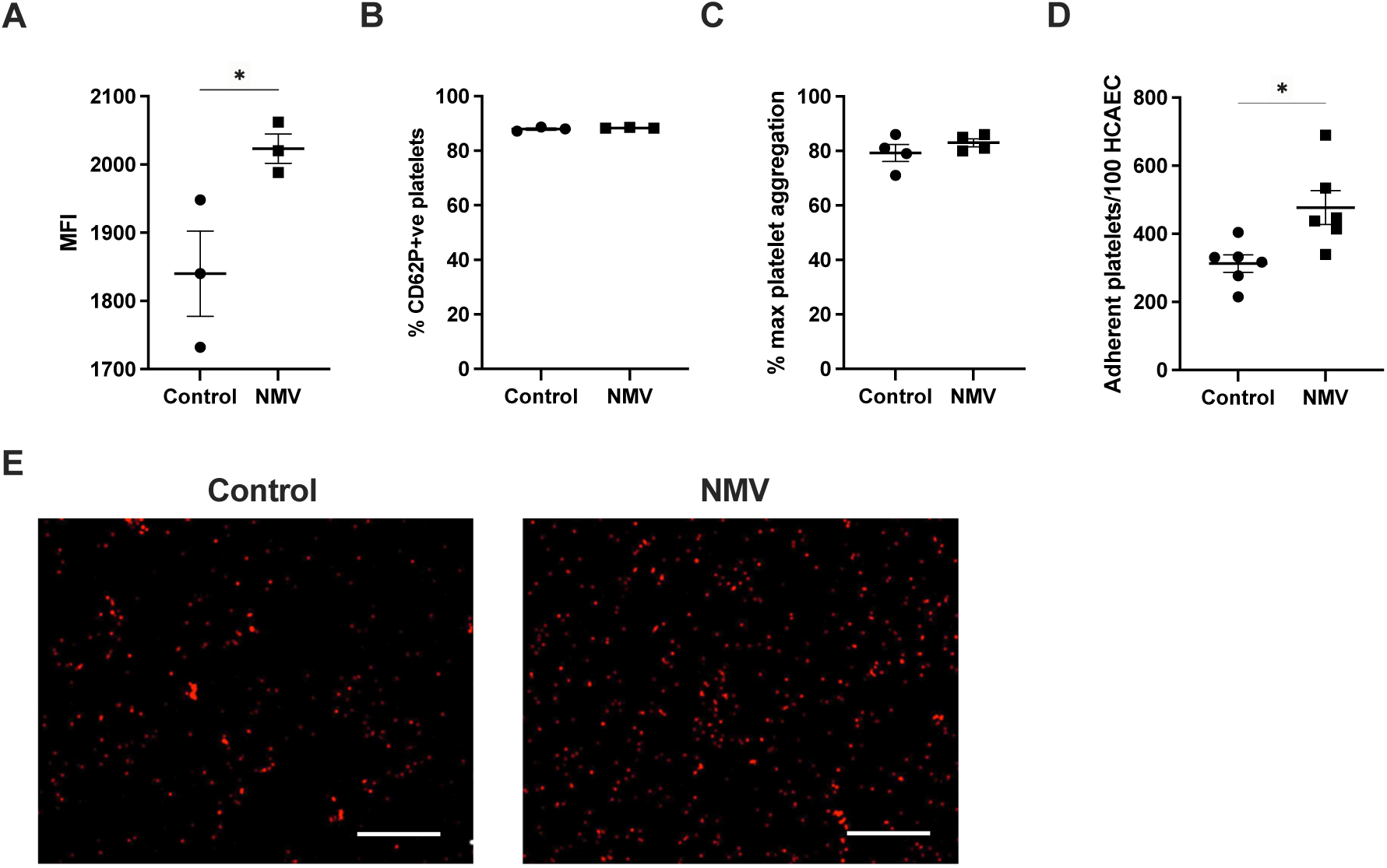
Platelet P-selectin expression, aggregation and adhesion to coronary artery endothelial cells in response to neutrophil microvesicles. Freshly prepared PRP was incubated with neutrophil microvesicles (NMVs) or supernatant control in combination with 5μM ADP prior to labelling with anti-CD42a and anti-CD62P antibodies. Fixed platelets were assessed for P-selectin expression by flow cytometry. **(A)** Median fluorescence intensity (MFI) of P-selectin and **(B)** percentage of P-selectin positive platelets in supernatant control and NMV treated platelets (n=3). To assess platelet aggregation, freshly prepared PRP was incubated with supernatant control or NMVs prior to addition of 0.25 μM Horm collagen. Platelet aggregation was assessed by light transmission aggregometry (LTA). **(C)** Percentage of maximal aggregation of platelets in NMV-treated compared to supernatant treated platelets (n=4). Human coronary artery endothelial cells (HCAECs) were cultured for 24 h prior to treatment with supernatant control or NMVs (1:5 ratio HCAEC:NMV). Cells were washed and incubated with PKH-26 labelled platelets at a 1:50 HCAEC:platelet ratio for 30 min. Cells were thoroughly washed to remove non-adherent platelets and platelet adhesion assessed by widefield fluorescence microscopy. (D) The number of adherent platelets per 100 HCEAC was determined using Image J analysis software (n=6). (E) Representative images of labelled platelets (red) adhered to HCAECs treated with supernatant control and NMVs. Scale bar = 100 µm. Data expressed as mean ±SEM. Statistical significance assessed by paired t-test. * p=≤0.05

## Discussion

Circulating MVs in the plasma are likely to be important mediators of many diseases and have been implicated in the pathogenesis of atherosclerosis ^18^. We have found that nLDL induces enhanced NMV release from neutrophils and that these NMVs contain active MMPs capable of degrading ECM proteins. This suggests that increased levels of NMVs in the circulation in response to elevated nLDL levels may increase the risk of plaque erosion. Characterisation of these NMVs demonstrated that they had a size range of 105-255 nm, expressed CD66b and had high expression of CD9 and undetectable expression of CD63. CD9 is a tetraspanin that is found in the plasma membrane, whereas CD63 is a marker of endosomal membrane derived EVs ^19^. These data show that the vesicles used in this study were derived from neutrophil plasma membrane rather than via the endosomal pathway associated with smaller EVs/exosomes.

MMP-8 and MMP-9 were the most abundant proteases in NMVs and differences in their relative abundance may be due to specific incorporation of these MMPs or merely reflect levels of the different proteases in the parent cell. These MMPs have previously been detected in NMVs from adherent neutrophils and neutrophils in suspension ^20^ and have been linked with plaque disruption ^14,21^. MMP-8 inactivation has been shown to limit lesion formation in ApoE^-/-^ mice and increase collagen content of plaques ^22^. Degradation of collagen and other ECM proteins is a key part in the hypothesised mechanism of plaque erosion making MMP-8 and MMP-9 relevant factors in this process. There is evidence to suggest MMP-9 also acts upon endothelial cells directly inducing apoptosis and inhibition of MMP-9 production in mice reduces atherosclerotic lesion size, indicating a role in plaque progression ^23^.

NMVs from unstimulated neutrophils contained 3 further proteases not detected in NMVs from nLDL stimulated neutrophils, namely urokinase (uPA), cathepsin X/Z/P and ADAM8. uPA has been previously reported in isolated neutrophils ^24^ and is interesting due to its role in thrombolysis, converting plasminogen to the thrombolytic protease plasmin ^25^ suggesting that uPA is protective against thrombosis. The lack of uPA in NMVs from nLDL stimulated neutrophils could indicate nLDL stimulation induces production of more pro-thrombotic NMVs. Cathepsin S and A were detected in both NMV populations whereas cathepsin X/Z/P was only detected in NMVs from unstimulated neutrophils. Cathepsin S has been previously linked with atherosclerosis ^26^ with cathepsin S knockout mice demonstrating reduced atherosclerosis as assessed by plaque size ^27^. Cathepsin X/Z/P has been reported to interact with integrins of ECs via its RGD-motif to support EC adhesion to the ECM ^28^. Cathepsin V was detected only within NMVs from nLDL stimulated neutrophils. Cathepsin V is a potent cysteine protease targeting the elastin components of the ECM and has been detected in human atheroma ^29^. Other studies have shown that whilst cathepsin K, S and V degrade elastin fibres and promote calcification, this process leads to protection of fibres from further degradation ^30^. Therefore, it is difficult to determine the exact effect of these cathepsins on promoting plaque erosion.

NE was also found to be contained within NMVs from both unstimulated and nLDL stimulated neutrophils. NE has been linked with EC dysfunction by inducing apoptosis via the PERK/CHOP pathway of the unfolded protein response (UPR) ^31^. Co-localisation of NE with apoptotic ECs in rupture-prone human plaques further supported the role NE may play in atherosclerosis ^31^. What is also important to plaque erosion is the effect of NE on the degradation of the ECM. It has been established for decades that NE degrades ECM proteins, collagens type I and type III ^32,33^ and more recently it has been shown to activate proteases, such as MMP-9 whilst also degrading inhibitors, such as TIMP-1 ^34^ therefore acting to exacerbate their action. This has been noted in cystic fibrosis (CF) where a higher MMP-9/TIMP-1 ratio was associated with NE and bronchiectasis ^35^.

Whilst the numbers of NMVs generated significantly increased following nLDL treatment, the concentration of MMP-9 and NE per NMV did not significantly differ, indicating that nLDL treatment did not lead to an enrichment of these proteins within NMVs. However, due to the significant increase in the number of NMVs generated following nLDL stimulation, the total amount of both MMP-9 and NE released from neutrophils and contained within NMVs was increased. Whilst NE activity was not higher in NMVs derived from nLDL stimulated neutrophils, the overall NE activity was significantly higher due to the increase in NMV released per neutrophil. MMP-9 activity was higher in NMVs from nLDL stimulated neutrophils and this activity was inhibited by TIMP-1, an irreversible inhibitor of a broad range of MMPs. The presence of active MMP-9 is important as it has the potential to degrade collagens I and III found in stable plaques ^16^ and can negatively affect EC integrity ^8^ which in combination may promote EC detachment from stable plaques.

Having established NMVs contain active proteases, it was determined that NMVs were able to degrade gelatin and that TIMP-1 significantly inhibited this activity. It was therefore hypothesised that NMVs may facilitate the detachment of HCAECs from ECM, a key contributing factor in plaque erosion. There is evidence to support this hypothesis; MMP-9 has been previously shown to degrade junctional proteins between epithelial cells and decrease epithelial cell integrity ^8^ and neutrophils have been shown to promote EC detachment in an in vitro model of plaque erosion ^5^. A higher proportion of HCAECs demonstrated a propensity to detach following treatment with NMVs compared with HCAECs treated with supernatant controls. This effect was largely inhibited by incubation with TIMP-1.

Whilst our data supported the hypothesis that degradation of the ECM by MMPs within NMVs promoted HCAEC detachment, it was important to investigate alternative mechanisms through which NMVs may be inducing HCAEC dysfunction and subsequent detachment. One potential mechanism was apoptosis. Durand *et al.* identified apoptosis as a relevant process in thrombosis demonstrating that induction of EC apoptosis in rabbit femoral arteries promoted endothelial detachment and thrombosis compared to saline treated control arteries ^36^ and this has since been replicated ^37^. Further studies have linked apoptosis specifically with plaque erosion; for example, Franck *et al.* showed via immunohistochemistry of human plaques that the rate of apoptosis is greater in erosion-prone plaques compared to rupture-prone plaques and further to this presented data correlating neutrophil adherence to erosion-prone plaques with this increased rate of EC apoptosis ^6^. In our study, the percentage increase in HCAEC apoptosis following NMV treatment rose by over 500% compared to supernatant controls. EC proliferation and migration in response to desquamation may help to prevent plaque erosion by maintaining an intact EC monolayer. NMVs were found to reduce EC proliferation and inhibited the wound healing response. These data are supported by previous work that has found NMVs inhibit wound healing by delivering myeloperoxidase ^38^.

Platelets are central to the process of thrombosis. In plaque erosion, where the cause of thrombosis is less apparent than in plaque rupture, subtle changes in platelet activation and propensity to adhere could provide useful insights into how thrombosis initiates and propagates. P-selectin is a widely used marker of platelet activation due to its presence on the platelet surface only following activation-induced degranulation ^17^. NMVs did not induce P-selectin expression in resting platelets but were able to increase the expression of P-selectin in ADP stimulated platelets. Previous studies have found expression of the counter receptor for P-selectin, P-selectin Glycoprotein

Ligand-1 (PSGL-1), on the surface of NMVs ^7^ and it is therefore possible that increased expression of P-selectin on platelets leads to an increase in the adhesion of NMVs leading to initiation of a positive feedback loop. Analysis of platelet adhesion to HCAEC demonstrated that NMVs induced a 50% increase in platelet adhesion. Taken together these data demonstrate that NMVs increase platelet activation and adhesion to ECs, key steps in thrombus formation.

Given the importance of endothelial cell removal in the context of plaque erosion, evidence presented here that NMVs promote extracellular matrix degradation, endothelial cell detachment and endothelial apoptosis suggest NMVs may contribute to endothelial cell loss. Our data also suggest NMVs reduce proliferation and the wound healing response in endothelial cells, potentially reducing the ability of the endothelium to recover, further promoting plaque erosion. Furthermore, NMVs increase platelet activation and adhesion to the endothelium, processes that contribute to thrombus formation. NMVs may therefore be a novel therapeutic target in the treatment of atherosclerotic plaque erosion.

